# Vascularizing neurospheroids to probe vascular contributions to α-synuclein pathology in Parkinson’s disease

**DOI:** 10.64898/2026.08.28.747883

**Authors:** Anika Alim, Sandra Lwin, Purnopama Saha, Yoongyeong Baek, Myungwoon Lee, Jungwook Paek

## Abstract

Neurodegenerative diseases are increasingly associated with vascular dysfunction beyond progressive neuronal degeneration, yet how vascular pathology contributes to disease progression remains poorly understood, largely due to the lack of a neurodegenerative disease model capable of capturing neuronal pathology alongside associated vascular dysfunction. Here, we developed a microengineered 3D vascularized brain tissue model that integrates neurospheroids with a self-assembled, perfusable vascular network to recapitulate key features of the neurovascular interface. Using this model, we investigated the vascular contribution to Parkinson’s disease pathology by introducing α-synuclein preformed fibrils into the engineered vasculature. Intravascular α-syn fibril exposure induced endothelial barrier disruption, vascular leakage, inflammation, and vascular regression. Notably, this vascular insult was accompanied by intraneuronal α-synuclein aggregation within neurospheroids, suggesting that vascular dysfunction may facilitate the exposure of neural tissue to pathogenic α-synuclein. Our neurodegenerative disease modeling approach establishes a versatile and tractable platform for investigating vascular contributions to neurodegenerative disease progression.

## Introduction

Neurodegenerative diseases, including Alzheimer’s disease, Parkinson’s disease (PD), and amyotrophic lateral sclerosis (ALS), represent a growing global health burden as life expectancy continues to increase^1–4^. Although each disease presents with distinct clinical and pathological features, these disorders are commonly defined by disease-specific proteinopathies that contribute to the progressive dysfunction and loss of vulnerable neuronal populations^5–7^. Increasing evidence, however, indicates that neurodegenerative pathology extends beyond neurons, with cerebrovascular dysfunction emerging as a shared pathological feature across this disease spectrum^8–12^. Patients with neurodegenerative diseases frequently exhibit blood-brain barrier (BBB) leakage, reduced cerebral blood flow, and microvascular abnormalities in brain regions affected by neurodegeneration^13–17^, raising the possibility that vascular impairment is not simply a downstream consequence of proteinopathy-associated neuronal damage but a critical contributor to disease progression. Despite growing recognition of its pathological relevance, the mechanisms by which vascular dysfunction shapes neurodegenerative disease progression remain incompletely understood.

In neurodegenerative diseases, pathogenic protein species such as amyloid-β, tau, and α-synuclein (α-syn) are not confined to the neuronal compartment. These species can be detected in cerebrospinal fluid and the bloodstream, where they may interact directly with endothelial cells and disrupt tight junction integrity and barrier function^18–20^. BBB compromise, in turn, may facilitate vascular-to-parenchymal transfer of pathogenic protein species, creating a feed-forward cycle in which barrier dysfunction promotes protein dissemination and regional neurodegeneration. Despite its potential relevance to neurodegenerative disease progression, direct experimental evidence for vascularly mediated protein dissemination remains limited. Current understanding relies largely on clinical observations, inference from the physiological roles of the cerebral vasculature, and experimental associations between amyloid pathology and BBB impairment. This gap reflects, in part, the difficulty of modeling vascular transport, endothelial barrier disruption, and neural exposure to pathogenic protein species within a single experimentally tractable system.

Existing models each capture only part of this process. Animal models provide physiological complexity but make it difficult to track intravascular protein transport in real time or isolate the specific contribution of the vasculature from other systemic and tissue-level factors^21,22^. Conventional in vitro BBB models provide useful readouts of endothelial barrier integrity and permeability^23–25^, but their reliance on porous membranes or thin-film substrates introduces an artificial support layer beneath the endothelium that may restrict or confound the direct passage of large amyloid fibrils into the neural compartment. As a result, these existing models are not well suited to directly examine how circulating pathogenic fibrils cross an impaired endothelial barrier and progressively expose otherwise healthy neural tissue.

Here, we demonstrate how synergistic integration of microengineering and vascularized tissue engineering can address a critical technical challenge in modeling vascular contributions to neurodegenerative disease progression. Leveraging human organ-on-a-chip technology, we created a 3D vascularized brain tissue model that recapitulates key features of the native human neurovascular interface (**Fig. 1a**) while enabling amyloid fibril transport through engineered vasculature and assessment of its pathological impact on neurons. Specifically, we leveraged a microengineered tissue culture platform established in our prior study^8^ to support the formation of a self-organized vascular network based on the vasculogenic principle. By incorporating organotypic neurospheroids within this vascular bed, we successfully constructed a vascularized neural tissue model that accurately recapitulates key structural and functional features of the neurovascular interface, including mature neurons with axonal and dendritic projections and perfusable blood vessels (**Fig. 1b**). Using Parkinson’s disease (PD) as a model disorder, we introduced α-syn fibril seeds into the vascularized neurospheroid model to reproduce PD-associated vascular pathology in vitro. This approach recapitulated key pathological vascular features, including endothelial barrier impairment, vascular inflammation, and vascular regression. In particular, intravascularly delivered α-syn fibrils seeded intraneuronal α-syn aggregation within neurospheroids, possibly through passage across the impaired endothelial barrier, a process that remains difficult to capture with existing experimental models (**Fig. 1c**). Beyond these disease-specific findings, this work establishes a generalizable strategy for engineering vascularized neural tissues and provides a platform for investigating a broader range of neurodegenerative and vascular brain diseases.

**Figure 1.**
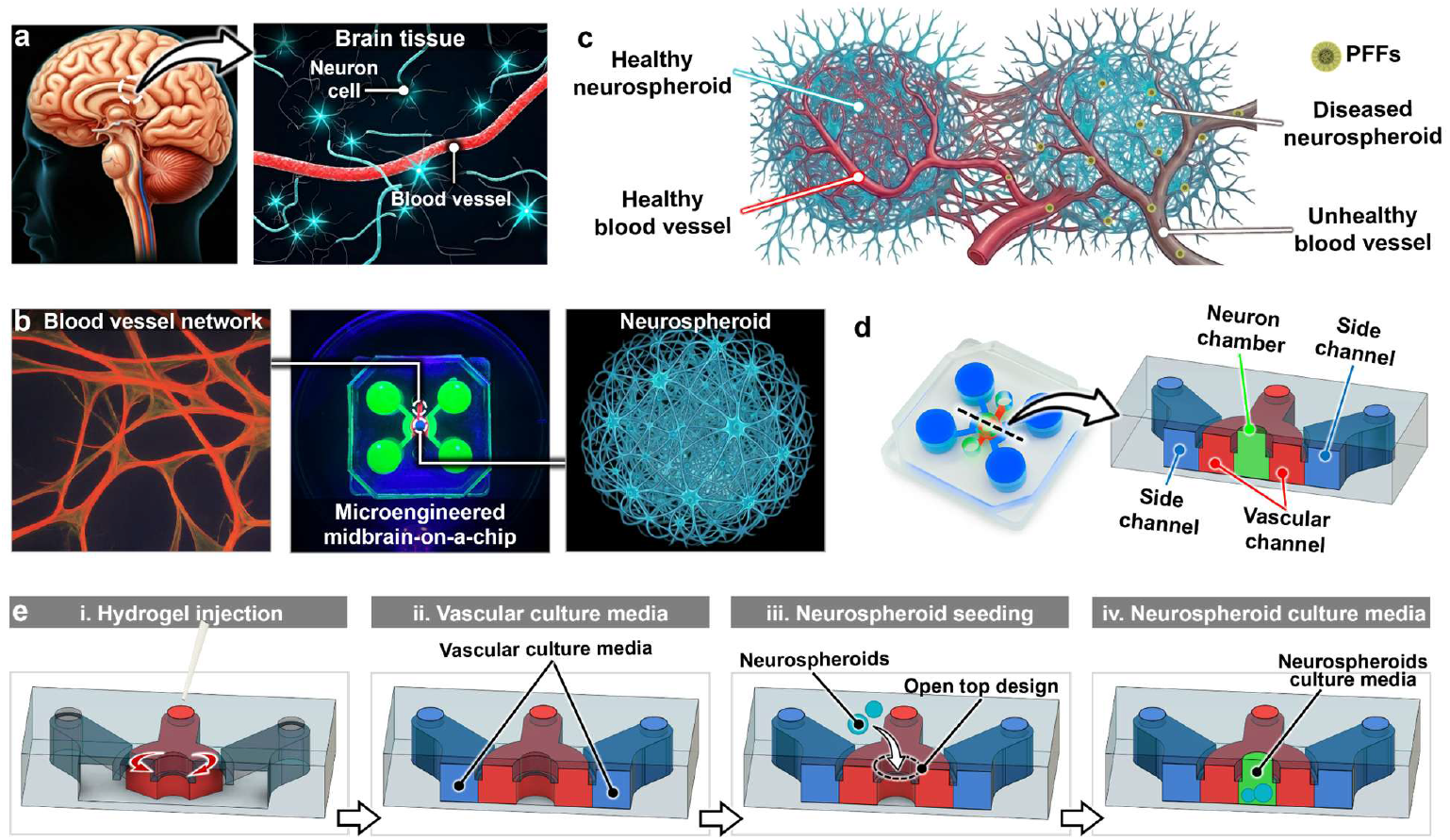
Microengineered human neurovascular model. **a)** Neurovascular interface in the human brain. **b)** Microengineered neurovascular model integrating a self-assembled vascular network with a neurospheroid. **c)** Schematic representation of PD-associated neuropathology and vascular dysfunction induced by α-synuclein preformed fibril (PFF) exposure in the vascularized neurospheroid model. **d)** Cross-sectional view of the microfabricated brain tissue culture platform showing the neuron chamber, vascular channel, and side channels. **e)** Schematic illustration of the sequential steps to construct the vascularized neurospheroid model. (i) Hydrogel solution containing vasculogenic cells is injected into the vascular channel. (ii) Following gelation, vascular culture medium is added to the side channels. (iii) After vascular network formation, neurospheroids are introduced into the neuron chamber through the open top. (iv) The neurospheroids and vascular network are cocultured to establish a vascularized neurospheroid model.

## Results and Discussion

### Device Design and Construction

The microphysiological model of vascularized brain tissue developed here builds upon a microfabricated culture platform established in our prior study^8^. A key feature of this platform is its multi-compartment design, which spatially organizes a 3D neuronal compartment within the central neuron chamber and a perfusable microvascular network within the flanking vascular channel, while allowing the two compartments to interface directly (**Fig. 1d**). Notably, this microengineering approach supports compartment-selective manipulation, each tissue component can be perturbed independently without disrupting the other, yet the two remain able to interact directly. For instance, vascular-directed stimuli introduced through the side channels perturb the engineered vasculature alone, isolating the effect of vascular dysfunction on the neuronal compartment, whereas the open-top neuron chamber allows neurospheroids to be manipulated directly to probe neuron-initiated effects on the surrounding vasculature. This bidirectional controllability positions the platform well for dissecting neurovascular interactions in neurodegenerative disease.

The platform was fabricated using standard soft-lithography with polydimethylsiloxane (PDMS), and vascularized neurospheroids were then constructed through the stepwise process schematically shown in **Fig. 1e**. Briefly, we first prepared a hydrogel solution containing vasculogenic cells and injected it into the vascular channel, where it remained in place without spilling into the neuron chamber or side channels due to capillary pinning at the thin gel-guides along the bottom surface (**Fig. 1e-i**). After thermal gelation, vasculogenic culture media was supplied through the side channels to support vascular network formation (**Fig. 1e-ii**). Once the vascular network was established, differentiated neurospheroids were seeded into the neuron chamber through the open top (**Fig. 1e-iii**). This open-top design allowed neurospheroids, or other cell types, to be incorporated without disrupting the neighboring preformed vasculature. Co-culture of the neurospheroids and vascular network then generated a 3D tissue construct in which neuronal tissue and engineered vasculature are structurally integrated and functionally coupled (**Fig. 1e-iv**).

### Engineering vascularized human brain tissue

The neuronal compartment of our vascularized brain tissue construct was reconstituted using neurospheroids, 3D organotypic neuron aggregates designed to recapitulate the complex structural integration with a surrounding vascular network that is a key anatomical feature of the native neurovascular tissue interface (**Figs. 1b, c**). Engineering of the brain construct therefore began with spheroid preparation, initiated by seeding human neuroblastoma cells into ultra-low attachment 96-well plates in growth media (**Fig. 2a, top-left**). Over the following days, cells progressively aggregated together and by day 12 post-seeding, they self-assembled into round, spherical spheroids (**Fig. 2a, top-right**). During spheroid formation, cells were sequentially exposed to two differentiation media formulations^26,27^, first at day 3 and again at day 8 post-seeding, to induce differentiation toward a mature neuronal phenotype (**Fig. 2a, bottom**). By day 12 post-seeding, following 10 days of differentiation, the spheroids retained their spherical architecture while exhibiting robust neuritic projections, a key structural hallmark of neuronal maturation (**Fig. 2b**). This neuronal differentiation was confirmed by immunofluorescent staining for microtubule-associated protein 2 (MAP2) and growth-associated protein 43 (GAP43), markers of dendrites and axons, respectively, which revealed complex neurite-like networks indicative of axonal and dendritic structures (**Fig. 2b, top-left**). β III-tubulin staining further confirmed the presence of an organized neuronal cytoskeleton (**Fig. 2b, top-right**), and the Live/Dead assay demonstrated high cell viability within neurospheroids (**Fig. 2b, bottom**).

**Figure 2.**
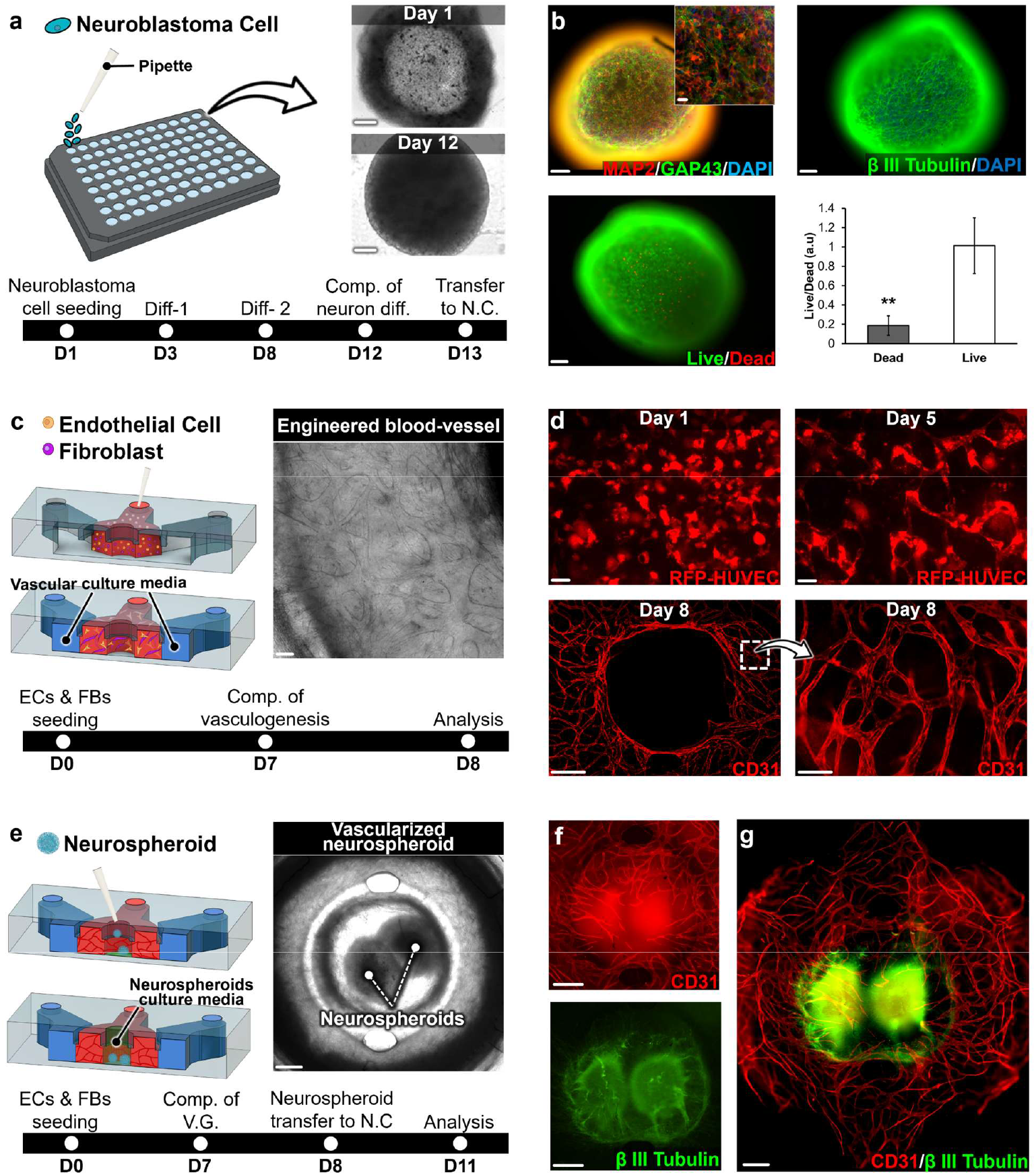
Construction of differentiated neurospheroids and their integration with engineered vasculature. **a)** Neurospheroid formation and differentiation begin with the seeding of neuroblastoma cells into ultra-low attachment microwells. During spheroid culture, the neuroblastoma cells undergo neuronal differentiation prior to transfer into the device, as indicated in the timeline. Abbreviations: D, day; N.C., neuron chamber; comp., completion; diff., differentiation. Bright-field images show morphological changes in neurospheroids on Days 1 and 12. Scale bars, 100 µm. **b)** Immunofluorescence staining of MAP2 (red) and GAP43 (green) shows dendritic and axonal structures, respectively. The inset shows a magnified view of MAP2 and GAP43 expression. βIII-tubulin staining identifies neuronal cytoskeletal structures, while Live/Dead staining confirms the viability of differentiated neurons within the spheroids. Scale bars, 100 µm and 25 µm for the inset. **c)** Schematic showing vasculogenic formation of engineered vessels within the device. A fibrin hydrogel containing endothelial cells and fibroblasts was introduced into the vascular channel, where the cells self-assembled into interconnected vascular networks over 8 days. A bright-field image of the engineered vascular network and the corresponding vasculogenic culture timeline are shown. Abbreviations: ECs, endothelial cells; FBs, fibroblasts. Scale bar, 100 µm. **d)** RFP-HUVECs co-cultured with fibroblasts progressively formed an interconnected endothelial tubular network from Day 1 to Day 5. CD31 immunostaining on Day 8 confirms the establishment of a complex vascular network within the vascular channel. Scale bars, 100 µm for the top-left, top-right, and bottom-right images and 500 µm for the bottom-left image. **e)** Schematic illustrating transfer of differentiated neurospheroids into the open-top neuron chamber for vascularization. The bright-field image shows neurospheroids integrated with the engineered vascular network, and the timeline summarizes the coculture of neurospheroids and vasculature within the device. Abbreviation: V.G., vasculogenesis. Scale bar, 500 µm. **f)** CD31 immunostaining shows engineered vessels extending from the vascular channel into the neuron chamber containing neurospheroids (β III-tubulin). Scale bars, 500 µm. **g)** Immunofluorescence analysis shows differentiated neurospheroids (βIII-tubulin) integrated with the surrounding CD31-positive vascular network. Scale bar, 1000 µm. ***P ≤ 0.001, **P ≤ 0.01, *P ≤ 0.05. Data show mean ± SD and P values were from unpaired, two-sided t-test.

In parallel with the neurospheroid preparation, we constructed a vasculogenic cell-laden hydrogel scaffold within the vascular channel to form complex 3D vascular networks emulating key structural features of native vasculature in the human brain. Briefly, a precursor fibrinogen solution was mixed with human umbilical vein endothelial cells (HUVECs) and normal human lung fibroblasts (NHLFs) and then supplemented with thrombin to initiate gelation. This cell-laden fibrin solution was then rapidly injected into the vascular channel prior to complete gelation, where it solidified into a stable fibrin hydrogel scaffold (**Fig. 2c, top-left**). Following gelation, the endothelial cells and fibroblasts were co-cultured with endothelial growth media to allow the formation of a vascular plexus within the fibrin gel (**Fig. 2c, top-right**) over 7 days (**Fig. 2c, bottom**), as described in our prior studies^8^. Time-lapse fluorescence imaging of RFP-HUVECs during this co-culture revealed the progressive development of vascular network structures within the gel (**Fig. 2d**). At early time points, endothelial cells began spreading throughout the hydrogel scaffold (**Day 1 in Fig. 2d**). HUVECs appeared significantly elongated and began forming chord-like structures over time and eventually led to the formation of cellular networks resembling vascular networks formed in vivo (**Day 5 in Fig. 2d**). Immunostaining on Day 8 further validated this vasculogenic process, showing interconnected networks of CD31-positive endothelial cells throughout the vascular channel (**Fig. 2d, bottom**). Notably, the central avascular region shown in the bottom-left microimage of **Fig. 2d** corresponds to the neuron chamber, which was intentionally left free of vasculature at this stage to subsequently introduce neurospheroids and enable their interface with the surrounding vascular network.

Once mature neurospheroids and a vascular bed were prepared, we next combined these two tissue components to generate vascularized neurospheroids within our microengineered device. To this end, differentiated neurospheroids were first suspended in an endothelial cell-laden collagen solution, which was then transferred into the neuron chamber through our device’s open top (**Fig. 2e, top-left**). Following the gelation of the cell-embedding collagen solution, the tri-culture of neurospheroids, endothelial cells, and the pre-formed vascular bed was then maintained for additional 3 days (**Fig. 2e, bottom**), during which endothelial cells within the collagen gel spread and organized into luminal structures that interconnected both with one another and with the adjacent engineered vessels of the vascular channel, forming a continuous vascular network surrounding the neurospheroids (**Fig. 2e, top-right**). Our immunostaining of CD31 demonstrated this angiogenic process in the neuron chamber by showing the presence of engineered vessels extended from the vascular channel to the neuron chamber containing neurospheroids (**Fig. 2f**). The successful tri-culture was also confirmed by immunostaining for our vascularized neurospheroids with CD31 labeling endothelial vessels and β III-tubulin identifying the mature neurons (**Fig. 2g**).

Importantly, this 3D tissue integration between the neuronal and vascular compartments enables faithful recapitulation of the physiological interactions between capillaries and brain parenchyma, which are central to understanding how vascular impairment contributes to neuronal dysfunction in neurodegenerative disease. The open-top design of the neuron chamber further addresses the challenge of distinct developmental timelines between neuronal and vascular tissue, permitting neurospheroids to be introduced into an already-established vascular bed without disrupting its structural integrity. Together, these capabilities demonstrate the broader applicability of our engineering approach beyond the present study, offering a versatile and tractable strategy for engineering vascularized neural tissues relevant to a range of vasculature-related brain diseases.

### Reconstituting α-syn-driven endothelial barrier dysfunction

Although vascular pathology is increasingly recognized as a key feature of neurodegenerative disease^28–30^, direct experimental evidence for its causal contribution to neurodegeneration remains scarce. Current understanding is largely indirect, drawn from clinical and post-mortem observations, reasoning based on established vascular physiology, and interpretations of experimental blood-brain barrier models. This knowledge gap reflects, in part, a longstanding emphasis on neuronal pathology in neurodegenerative disease research, which has left the accompanying vascular dysfunction comparatively overlooked and limited efforts to model neurodegeneration-associated vascular pathology. The lack of physiologically faithful models has, in turn, constrained mechanistic insight into how vascular impairment influences disease progression. To address this gap, we applied our vascularized neurospheroid model to examine how vascular dysfunction contributes to proteinopathy, a hallmark of many neurodegenerative diseases. Using PD as a model disease, we specifically investigated how endothelial barrier impairment contributes to the spread of blood-borne α-syn fibrils and their subsequent seeding of intraneuronal α-syn aggregation, a defining pathological hallmark of PD^31,32^.

Probing this vascular contribution, however, required a perfusable engineered vasculature capable of emulating intravascular delivery of α-syn fibrils and their passage across the endothelial barrier. To this end, endothelial cells were introduced into the two side channels of the device, where they structurally integrated with the engineered vessels formed in the vascular channel (**Fig. 3a**). This anastomosis-like process established a continuous luminal pathway connecting the side channels with the vascular network (**Fig. 3a, bottom**), allowing fluid to flow through the engineered vessels and thereby recapitulating perfusability, an intrinsic functional property of in vivo blood vessels^8,33^. This vascular functionality was demonstrated by introducing a 40 kDa fluorescein isothiocyanate (FITC)-dextran solution into one side channel and tracking its flow through the tree-like branched network of engineered vessels into the neuron chamber (**Fig. 3b**). Over 5 min, the tracer remained confined to the vascular regions, with no significant change in fluorescence intensity in the extravascular area, indicating endothelial barrier integrity (**Fig. 3b**). An overlay image of CD31-positive vessels and FITC-dextran flow further confirmed both the perfusability of the engineered vasculature and its endothelial barrier function (**Fig. 3c**).

**Figure 3.**
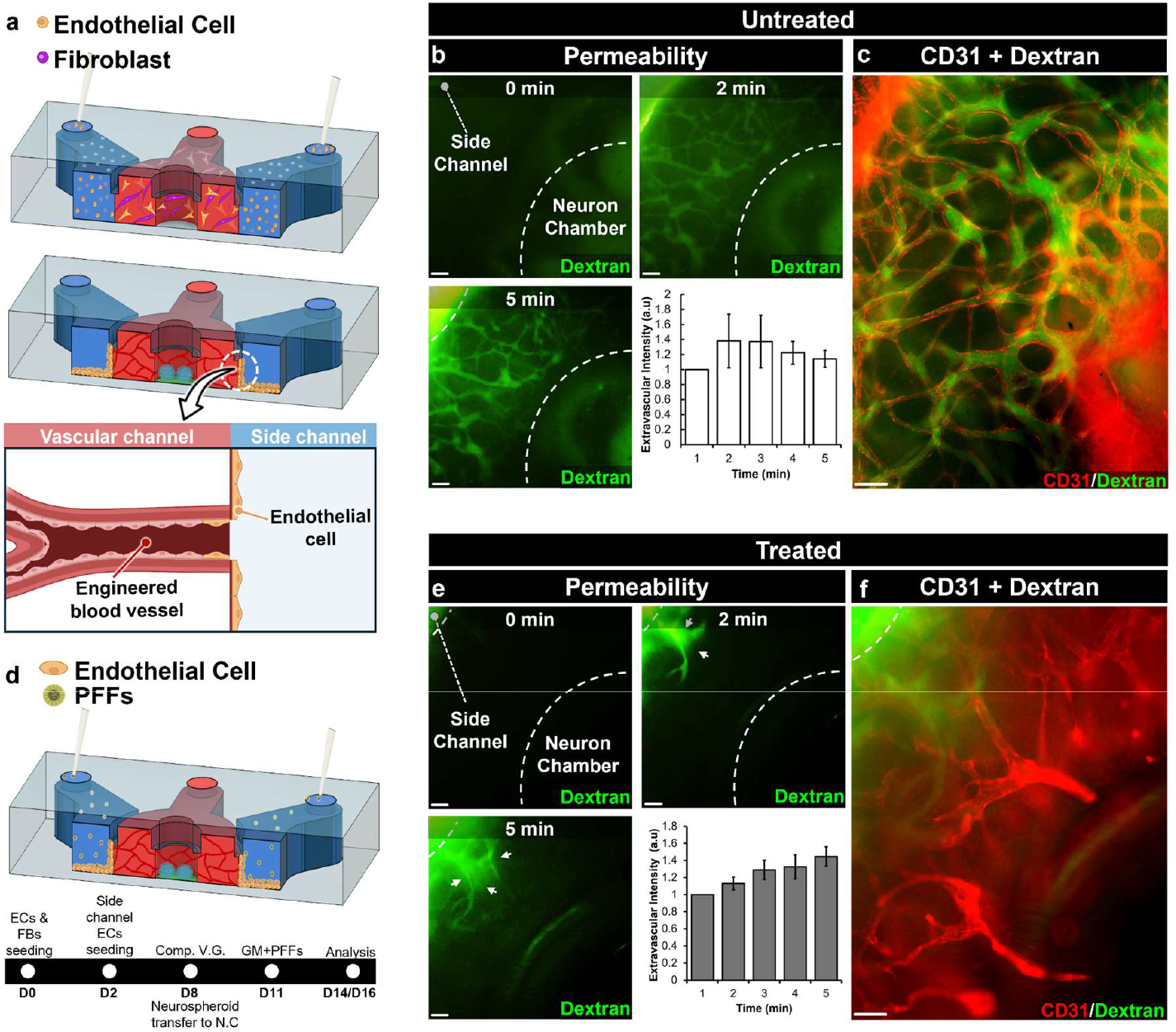
Microphysiological modeling of PD-associated endothelial barrier disruption. **a)** Schematic illustration of the formation of a perfusable vascular network through anastomosis between endothelial cells in the vascular channel and those seeded in the two side channels. **b)** FITC-dextran perfusion in untreated devices. The fluorescent tracer remains largely confined within the intravascular region over 5 min. Quantification of extravascular dextran fluorescence intensity shows no progressive increase over time. Scale bar, 100 µm. **c)** Merged fluorescence image showing FITC-dextran (green) confined within the CD31-positive vascular network (red). Scale bar, 100 µm. **d)** Schematic illustration of α-syn PFF treatment for 3 days followed by 3 days of culture in fresh medium prior to analysis. Abbreviations: ECs, endothelial cells; FBs, fibroblasts; V.G., vasculogenesis; comp., completion; N.C., neuron chamber; G.M., growth medium; PFFs, preformed fibrils; D, day. **e)** FITC-dextran perfusion in α-syn PFF-treated devices shows dextran leakage into the extravascular region within 2 min (white arrows), followed by a further increase in extravascular fluorescence by 5 min. Quantification confirms a progressive increase in extravascular dextran fluorescence intensity over time. Scale bar, 100 µm. **f)** Merged fluorescence image showing FITC-dextran leakage (green) from CD31-positive vessels (red). Scale bar, 100 µm. ***P ≤ 0.001, **P ≤ 0.01, *P ≤ 0.05. Data show mean ± SD and P values were from unpaired, two-sided t-test.

Having established a perfusable vasculature, we next investigated whether our approach could recapitulate a disease-relevant vascular phenotype associated with PD. Because α-syn fibril exposure has been linked to endothelial barrier disruption, we asked whether this phenotype could be reproduced by using α-syn preformed fibrils (α-syn PFFs), short fibrillar assemblies widely adopted to induce both α-syn-driven neuropathology and blood-brain barrier dysfunction in PD-relevant experimental models^8,34–36^. To this end, α-syn PFFs were prepared as described in our prior study^8^ and introduced into the two side channels, from which they flowed into the engineered vessels in the vascular channel (**Fig. 3d**). Devices were treated with α-syn PFFs at 5 µM, a concentration selected based on our prior study^8^. After 3 days of treatment, fresh media was supplied and cultures were maintained for an additional 3 days prior to analysis (**Fig. 3d, bottom**). We then assessed whether α-syn PFF treatment compromised the endothelial barrier function of the engineered vessels, again using 40 kDa FITC-dextran. As shown in the time lapse micro images (**Fig. 3e**), α-syn PFF-treated vessels exhibited pronounced FITC-dextran leakage into the extravascular space within 2 min of perfusion (white arrows) with fluorescence intensity increasing further by 5 min, indicating loss of endothelial barrier integrity. Consistent with this observation, quantification of extravascular fluorescence intensity showed a gradual increasing trend over time following α-syn PFF treatment (**Fig. 3e**). It should be also noted that α-syn PFF treatment also compromised the structural integrity of the vasculature (**Fig. 3f**), reducing the density of perfusable vessels within the device (**Fig. 3e, f**).

Collectively, our approach reconstitutes not only the anatomical organization of in vivo vascular networks but also key physiological functions, including intravascular biomolecular transport and endothelial barrier function. This combination of structural and functional vascular features is essential for modeling how amyloid fibrils move through the vasculature, interact with the endothelial barrier, and ultimately gain access to the surrounding neural tissue. The platform therefore enables direct investigation of vascularly mediated fibril dissemination and of progressive fibril exposure to otherwise healthy neurons, providing a new experimental framework for examining how neurodegenerative pathology may propagate across anatomically distinct brain regions, an investigation that remains difficult with existing experimental systems^21–23^.

In addition to barrier disruption, α-syn PFF treatment also compromised vascular architecture, leading to a marked reduction in the density of perfusable vessels within the device (**Fig. 3f**). This structural deterioration is consistent with microvascular regression reported in PD patients and further supports the ability of the model to recapitulate PD-associated vascular impairment^37–39^. Importantly, the ability to capture both endothelial barrier failure and loss of perfusable vascular networks provides an opportunity to experimentally examine how a broad spectrum of vascular pathologies may influence downstream neurodegenerative processes.

Despite these unique capabilities, a limitation of our study is the use of HUVECs rather than brain-derived endothelial cells, which may constrain the physiological relevance and translational applicability of the model. HUVECs have nonetheless been widely used as a practical endothelial source for modeling BBB function and its pathological disruption in neurodegenerative disease, owing to their capacity to reproduce key BBB-like properties across both in vitro and in vivo settings^40–44^. Consistent with this precedent, our engineered vessels exhibited functional barrier integrity, restricting the extravascular diffusion of a 40 kDa fluorescent dextran, a tracer commonly used to assess BBB permeability^45–47^. This observation supports the use of HUVEC-based vascular networks as a practical basis for recapitulating BBB-like barrier function in our model, while also indicating a clear opportunity to improve physiological relevance in future studies by incorporating brain-derived endothelial cells.

### Modeling vascular contribution to α-syn pathology

Building upon our platform’s ability to model α-syn PFF-driven impairment of endothelial barrier integrity, we next asked whether the fibril-induced barrier disruption enables intravascularly delivered α-syn PFFs to reach neurospheroids in the neuron chamber and seed intraneuronal α-syn aggregation, a defining pathological feature of PD^48–50^. To test this, α-syn PFFs were introduced into the side channels at 5 µM and maintained for 3 days as described in the previous section, allowing fibrils to enter the engineered vascular network and compromise the endothelial barrier. We then assessed whether the fibrils had crossed the impaired barrier, reached the neuron chamber, and interacted with neurons, by immunostaining for intraneuronal α-syn aggregation within the neurospheroids.

For this investigation, we first established the untreated baseline to determine whether α-syn aggregation emerged spontaneously in the absence of fibril exposure. As shown in **Fig. 4a**, untreated devices exhibited no discernible intraneuronal α-syn aggregation, which typically presents as punctate structures within neurons^51,52^. In contrast, α-syn PFF treatment produced α-syn aggregation within the neurospheroids, evidenced by the appearance of aggregation-positive puncta (**Fig. 4a**). Although further validation is required to directly track fibril translocation across the endothelial barrier into the neuron chamber, these results suggest that α-syn PFF-induced barrier impairment permits vascularly delivered fibrils to access the neuronal compartment and template intraneuronal α-syn aggregation. Notably, this vascular transfer is unlikely to reflect passive permeability through an intact barrier, as α-syn PFFs are supramolecular fibrillar assemblies that are substantially larger than soluble 40 kDa dextran tracer commonly used to assess BBB permeability^53^.

**Figure 4.**
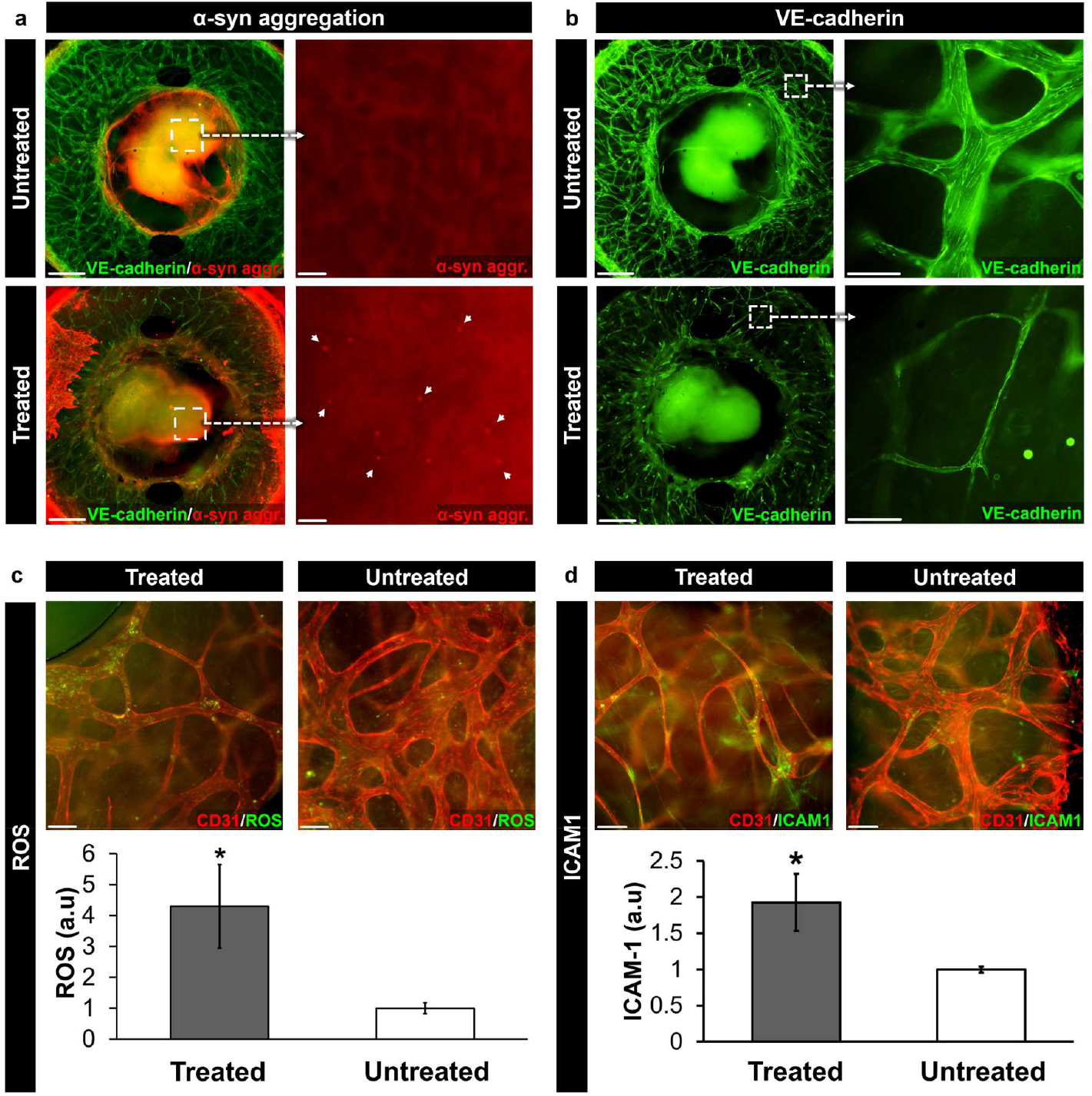
Investigation of vascular contributions to α-syn pathology in the vascularized neurospheroid model. **a)** Immunofluorescence analysis of α-syn aggregation shows punctate pathological α-syn accumulation within neurospheroids following α-syn PFF introduction exclusively through the side channels, whereas punctate aggregates are not observed in untreated controls. Scale bars, 500 µm for the left images and 25 µm for the right images. **b)** VE-cadherin immunofluorescence shows disrupted endothelial junctional organization following treatment. Treated devices exhibit fragmented vascular structures compared with untreated controls. Scale bars, 500 µm and 100 µm. **c)** α-syn PFF treatment induces oxidative stress within engineered vascular networks. ROS quantification shows a significant increase in the treated group compared with untreated controls. Scale bar, 100 µm. **d)** ICAM-1 immunofluorescence shows increased endothelial inflammatory activation in treated vascular networks relative to controls. Quantification confirms increased ICAM-1 expression following treatment. Scale bar, 100 µm. ***P ≤ 0.001, **P ≤ 0.01, *P ≤ 0.05. Data show mean ± SD and P values were from unpaired, two-sided t-test.

Together with the barrier leakage demonstrated earlier (**Fig. 3e**), the emergence of intraneuronal α-syn aggregation following intravascular PFF delivery further supports that α-syn PFF treatment compromised endothelial barrier integrity. To determine whether this functional compromise involved disruption of endothelial cell-cell junctions, critical molecular components for maintaining barrier function, we assessed the expression of VE-cadherin, a key adherens junction protein^54^. Consistent with the functional barrier impairment (**Fig. 3e**), VE-cadherin immunostaining revealed reduced signal intensity in α-syn PFF-treated devices compared with untreated controls, indicating disruption of endothelial adherens junctions (**Fig. 4b**). Notably, treated devices also exhibited an increased number of thin, acellular vascular structures resembling string vessels (**Fig. 4b**), a pathological vascular feature reported in PD patients^55,56^.

Having observed both barrier disruption and vascular structural deterioration, we next examined whether our modeling approach could capture additional PD-associated vascular pathologies. To this end, we focused on endothelial oxidative stress and inflammatory activation because these processes are key features of vascular dysfunction and may contribute to both barrier impairment and vascular regression^57– 59^. Accordingly, we assessed reactive oxygen species (ROS) production and intercellular adhesion molecule-1 (ICAM-1) expression, with ROS serving as a major mediator of oxidative endothelial injury^60,61^ and ICAM-1 reflecting inflammatory endothelial activation^62^. Consequently, our fluorogenic and immunofluorescence analyses revealed that α-syn PFF treatment significantly increased ROS levels and ICAM-1 expression compared with untreated controls (**Figs. 4c, d**). Together, these findings demonstrate that our approach can model a broad spectrum of vascular pathologies, including inflammatory endothelial responses that may accompany and contribute to barrier impairment and vascular regression during PD pathogenesis.

In summary, these results demonstrate that our vascularized neurospheroid model successfully recapitulates a cascade of PD-associated vascular pathologies driven by α-syn PFFs. Notably, intravascularly delivered fibrils seeded intraneuronal α-syn aggregation within the neurospheroids, probably by crossing the impaired endothelial barrier, a process difficult to capture with existing experimental models. Although further study is required, this finding provides experimental support for a vascular contribution to the spread of neurodegenerative pathology. α-syn PFF treatment further elicited endothelial oxidative stress and inflammatory activation, together with the emergence of string vessel-like vascular structures. Together, these observations establish that our model captures multiple interconnected features of PD-associated vascular impairment, spanning barrier breakdown, junctional loss, inflammatory activation, and structural regression

## Methods

### Device fabrication

To fabricate the device standard soft-lithography techniques were used. The top part of our device consisted of a poly(dimethylsiloxane) (PDMS) slab patterned with recessed microchannels and reservoirs on both its top and bottom surfaces. We also produced a plain PDMS slab following the same microfabrication process which served as the bottom part. The cross-sectional dimensions of the neuron chamber, vascular channel, and side channels were 2 mm (width) × 700 μm (height), 1 mm (width) × 700 μm (height), and 2 mm (width) × 700 μm (height), respectively (**Supplementary Fig.1**). The diameters of the opening in the neuron chamber and each were 2 mm and 5 mm, respectively (**Supplementary Fig.1**). For fabrication of these components, PDMS (Sylgard 184, Dow Corning) base and curing agent were thoroughly mixed at a weight ratio of 10:1 (base:curing agent) and poured onto 3D-printed masters (Protolabs). After degassing, PDMS was fully cured in an oven maintained at 55°C. The hardened PDMS was then carefully peeled from the molds, and inlet and outlet ports were punched at both ends of the vascular channel and the two side channels in the microstructured slab to provide fluidic access. For device assembly, the microstructured slab was stamped onto a thin layer of uncured PDMS, prepared by spin-coating at 2500 rpm for 2 min, and sealed against the plain PDMS slab that was used as the bottom part of the device. The assembled device was then baked at 55 °C to fully cure the PDMS adhesive.

Before cell culture, the assembled device was first sterilized by autoclaving with high-temperature pressurized steam (EZ Plus, Tuttnauer) for 1 hour. Following sterilization, a 2 mg/mL polydopamine (PDA) solution (Sigma, H8502) prepared in 10 mM Tris-HCl (pH 8.5) was introduced into the neuron chamber, vascular channel, and side channels. This PDA coating enhanced extracellular matrix (ECM) hydrogel and cell adhesion to the PDMS surfaces within the device. After incubation at room temperature (RT) for 2 hours, the PDA solution was gently aspirated, and the PDMS surfaces were washed twice with sterile-filtered deionized (DI) water to remove any unbound PDA molecules. The washed device was kept sterile at RT until use.

### Cell culture

To generate 3D neuronal cell cluster within our device, we used the human neuroblastoma cell (SH-SY5Y; ATCC, CRL-2266) and expanded using 25 cm^2^ flasks according to the manufacturer’s protocols. To culture SH-SY5Y cells, Dulbecco’s Modified Eagle Medium (DMEM; 10-017-CV, Corning) was used as the basal growth medium, supplemented with 10% (v/v) heat-inactivated fetal bovine serum (hiFBS; SH30396.03, Cytiva), 1% (v/v) GlutaMAX-I (35050-06, Thermo Fisher) and 1% (v/v) penicillin-streptomycin (SV30010, Cytiva). For device culture, SH-SY5Y cells at passages 3 to 10 were, as recommended by the cell supplier.

To establish 3D vascular network in our device, we utilized primary human umbilical vein endothelial cells (HUVECs; C2519A, Lonza) and primary normal human lung fibroblasts (NHLFs; CC-2512, Lonza). HUVECs and NHLFs were cultured in 75 cm^2^ flasks following the manufacturer’s protocols. These cells were cultured and maintained using endothelial cell growth medium (EGM)-2 (CC-3162, Lonza) and fibroblast growth medium (FGM)-2 (CC-3132, Lonza), media respectively. HUVECs and NHLFs between passage 2 and 5 were used for device culture.

### Neurospheroid generation

For neurospheroid formation, SH-SY5Y cells were trypsinized and resuspended in their growth medium and seeded them into ultra-low attachment 96-well plates at a density of 1 × 10^6^ cells/mL. The initial culture was maintained in their growth medium to promote neurospheroid formation. Two days after seeding, the SH-SY5Y cell growth medium was replaced with the first differentiation medium consisting of DMEM supplemented with 2.5% (v/v) hiFBS, 1% (v/v) GlutaMAX-I, 1% (v/v) penicillin-streptomycin, and 10 μM retinoic acid (50-165-6969, Fisher Scientific) to induce differentiation into dopaminergic neuronal cells. After 5 days, the first differentiation medium was switched to a second differentiation medium based on Neurobasal-A (10888022, Thermo Fisher) supplemented with 50 ng/mL brain-derived neurotrophic factor (BDNF; P23560, FUJIFILM Irvine Scientific), 20mM potassium chloride (KCl; 7447-40-7, Avantor Science), 1% (v/v) B27 (17-504-044, Fisher Scientific), 1% (v/v) GlutaMAX-I, and 1% (v/v) penicillin-streptomycin. The second differentiation medium was also maintained for an additional five days. Both the first and second differentiation media were refreshed every other day to support the maturation of neurospheroids.

### Vasculogenic cell culture and neurovascular co-culture

To form a vascularized tissue scaffold in our device, trypsinized HUVECs and NHLFs were resuspended in their growth media and mixed with fibrinogen (10 mg/mL; F8630, Sigma), thrombin (2 U/mL; T7513, Sigma), and aprotinin (1 U/mL; A1153, Sigma). The final density of HUVECs and NHLFs were 3.5 × 10^6^ cells/mL. 20 μL hydrogel solution containing the cells was injected into the vascular channel. The device was then incubated at 37°C with 5% CO_2_ for 30 min to allow thermal gelation. Once the fibrin gel solidified, EGM-2 was introduced into the top medium reservoirs and the side channels to support vasculogenesis.

Following vascular network formation, differentiated neurospheroids were carefully transferred into an endothelial cell-laden collagen solution containing 2 × 10^6^ cells/mL endothelial cells and 2 mg/mL collagen. The neurospheroids-collagen mixture was then injected into the neuron chamber through its open-top. After collagen gelation, the neurospheroids and endothelial cells were co-cultured for 3 days in a mixture of endothelial growth medium and secondary neurospheroid differentiation medium to support vascularization of the neurospheroids within the neuron chamber.

### Viability Assay

To assess the cell viability within the differentiated neurospheroids we performed Live/Dead assay using Live/Dead Cell Imaging Kit (488/570, ThermoFisher) according to the manufacturer’s protocol. The differentiated neurospheroids were incubated with the live-dead cell dye at room temperature for 15 min while protected from light. Subsequently, the neurospheroids were washed with DPBS to remove excess dye and imaged using an inverted fluorescence microscope (Eclipse Ti2, Nikon). Live cells exhibited green fluorescence, whereas dead cells were identified by red fluorescence.

### Transmission electron microscopy of α-syn PFFs

Negatively stained α-syn PFFs were prepared on 300-mesh lacey carbon coated copper grids (Electron Microscopy Sciences, Ultrathin Carbon Films, LC-325-CU-CC). A 10 μL aliquot of the PFF solution was deposited onto the grids and allowed to absorb for 1–2 min. Subsequently, the excessive solution was blotted, followed by a wash with 10 μL of water and then blotted again and stained with 10 μL of UranyLess EM stain solution (Electron Microscopy Sciences) for 30 seconds. TEM images were acquired using a JEOL 2100 F Field-emission transmission electron microscope at 120 kV, equipped with a Gatan Ultrascan CCD camera. The obtained images were processed using Digital Micrograph (GMS3) software (Gatan Inc.).

### Treatment of α-syn PFFs

To investigate the pathological effects of α-synuclein fibrils, the vascularized neurospheroid model was exposed to α-syn PFFs. For treatment, the stock solution of α-syn PFFs was first sonicated for 15-20 sec to ensure their uniform distribution and optimal working size (∼100 nm). The sonicated PFF stock was then diluted in EGM-2 medium to a final concentration of 5 μg/mL to assess its effects on the vascular component. The PFF-containing medium was introduced into the side channels, and the device was incubated for 72 h. Following treatment, the PFF-containing medium was replaced with fresh medium in both the neuron chamber and side channels. The cultures were then maintained for an additional 3 days before analysis.

### Measurement of Reactive Oxygen Species (ROS) production

Following the treatment with the PD-associated α-syn PFFs, the CellROX Green (5 μM in DPBS; C10444, ThermoFisher) were used to measure oxidative stress of the vascular network in our device. The dye solutions were injected into the device through the neuronal chamber and the two side channels and incubated at 37°C for 30 min. After incubation, the device was washed three times using DPBS and the vessels were imaged using an inverted fluorescence microscope (Eclipse Ti2, Nikon). To quantitively assess the data, fluorescent intensity was averaged from 3 devices for each experimental condition.

### Immunostaining and Quantification

For immunostaining, cells within our device were fixed with 4% paraformaldehyde (J19943-K2, ThermoFisher Scientific) for 30 min at RT and washed twice using DPBS. After fixation, they were permeabilized with 0.1% Triton X-100 (9002-93-1, VWR) for 15 min. To prevent non-specific binding, the cells were blocked with 1.5% bovine serum albumin (BSA; 22013, Biotium) for 30 min at RT. Subsequently, the cells were incubated overnight at 4°C with primary antibodies that selectively bind to the proteins of interest.

For imaging of the self-assembled vessels, we used a rabbit polyclonal anti-CD31 antibody (ab28364, 1:300-1:500, Abcam). To visualize the neuronal differentiation of neurospheroid was demonstrated using primary antibodies of rabbit polyclonal anti-MAP2 (A16829, 1:200, Antibodies.com) for dendritic extensions, mouse monoclonal anti-GAP43 (A85392, 1:1000, Antibodies.com) for axonal projections, and mouse monoclonal anti-β III Tubulin (A86691, 1:500, Antibodies.com) for the neuronal cytoskeleton. To confirm α-synuclein aggregation, anti α-synuclein aggregate antibody [MJFR-14-6-4-2] (A209538, 1:5000, abcam). For investigating inflammatory responses, mouse monoclonal anti-ICAM-1 (A85684, 1:500, Antibodies.com) antibodies. For analyzing vascular endothelial integrity, we used a mouse monoclonal anti VE-cadherin antibody (14-1449-82, 1:100, ThermoFisher).

After incubation with primary antibodies, the cells were washed twice with DPBS and incubated for 1-2 hours at RT with fluorescently labeled secondary antibodies (A32732, ThermoFisher; A32723, ThermoFisher; A32731, ThermoFisher; A11032, ThermoFisher). We also used Hoechst (33342, ThermoFisher) for nuclear staining.

Finally, fluorescence images of the cells were captured using the Nikon Ti2-Eclipse microscope, and image processing was performed using NIS Elements software.

### Testing of vascular perfusability

To investigate the perfusability of the engineered blood vessels, we used 40 kDa FITC dextran (50 μg/mL in PBS; FD40S-100MG, 60842-46-8, Sigma). The dextran solution was introduced through the side channels, a gradient of hydrostatic pressures across the hydrogel scaffold drove the fluorescent tracers through the vessels. Vascular perfusion was monitored and visualized using Nikon Ti2-Eclipse microscope and NIS-Elements. Time lapse images were captured over 5 min to track the movement of dextran within the vessels.

To generate quantitative data, extravascular fluorescence intensity was measured from four regions of interest per device at each time point. The intensity measured in each region was normalized to its corresponding baseline intensity at the initial time point, and the normalized values were then averaged across the four regions for each time point. Three devices were analyzed for each control and experimental group.

### Statistical analysis

Sample sizes for each experiment were based on a minimum of *n* = 3 independent devices per experimental group. Data was assessed using unpaired two-tailed t-tests and are presented as mean ± SD.

## Acknowledgements

We thank H Park for his input. This work was supported by the National Institutes of Health (NIH) (grant no. 1R21NS139178-01); Binghamton University (grant nos. TAE 1182867, ADLG329).

## Author contributions

A.A. and J.P. designed the research, performed the experiments, analyzed the data and wrote the manuscript. S.L. and P.S. helped with the experiments. Y.B. and M.L. produced short form α-synuclein fibrils and wrote the manuscript.

---

**Supplementary Figure 1:**
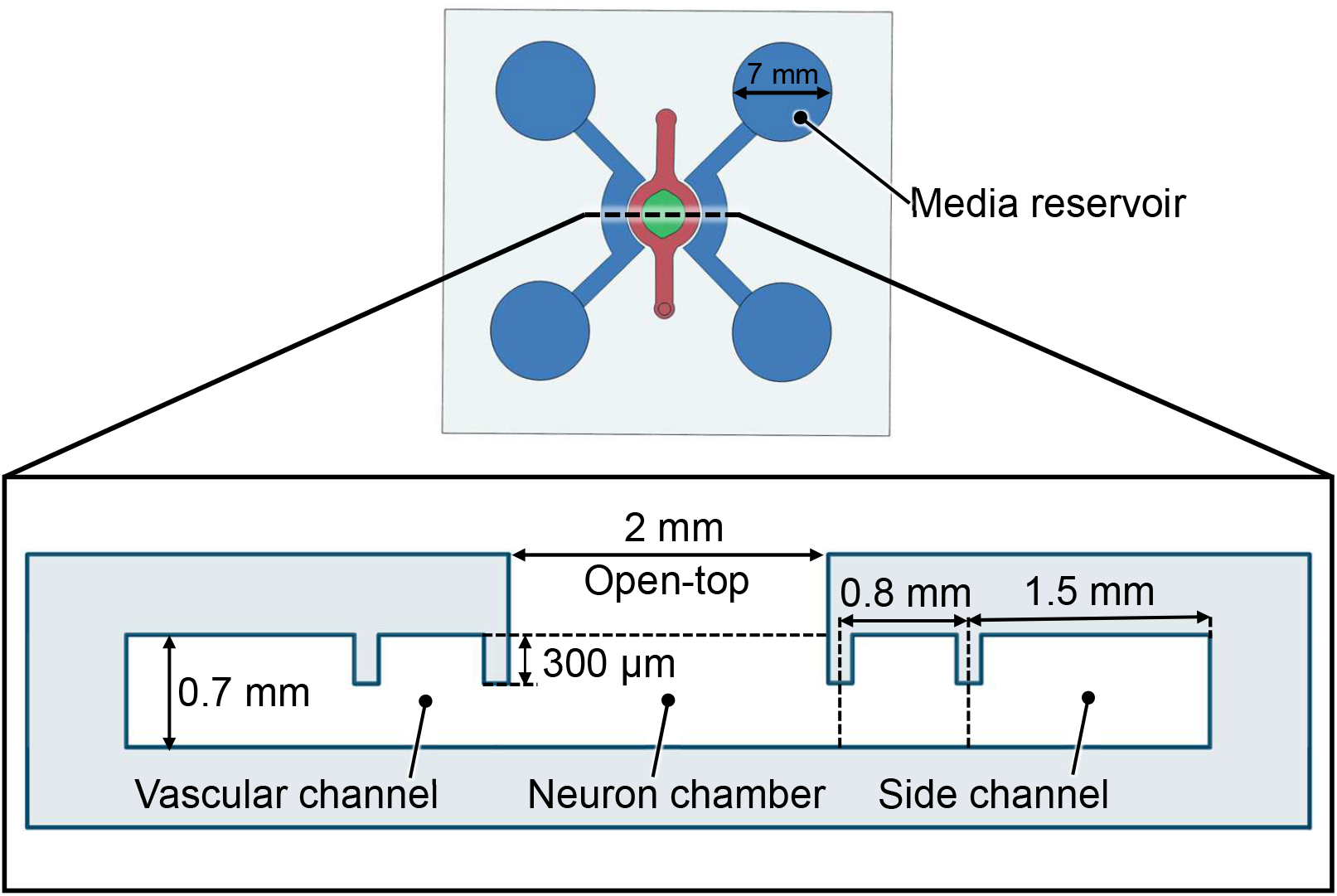
The diameter of each media reservoir is 7 mm and the cross-sectional dimensions of the neuron chamber, vascular channel, and side channels are 2.0 mm (width) × 0.7 mm (height), 0.8 mm (width) × 0.7 mm(height), and 1.5 mm (width) × 0.7 mm (height), respectively.

